# BALLI: Bartlett-Adjusted Likelihood-based LInear Model Approach for Identifying Differentially Expressed Gene with RNA-seq Data

**DOI:** 10.1101/344929

**Authors:** Kyungtaek Park, Jaehoon An, Jungsoo Gim, Sungho Won

## Abstract

**Motivation:** Transcriptomic profiles can improve our understanding of the phenotypic molecular basis of biological research, and many statistical methods have been proposed to identify differentially expressed genes under two or more conditions with RNA-seq data. However, statistical analyses with RNA-seq data often suffer from small sample sizes, and global variance estimates of RNA expression levels have been utilized as prior distributions for gene-specific variance estimates, making it difficult to generalize the methods to more complicated settings. We herein proposed a <u>B</u>artlett-<u>A</u>djusted <u>L</u>ikelihood based <u>LI</u>near mixed model approach (BALLI) to analyze more complicated RNA-seq data. The proposed method estimates the technical and biological variances with a linear mixed effect model, with and without adjusting small sample bias using Bartlett’s corrections.

**Results:** We conducted extensive simulations to compare the performance of BALLI with those of existing approaches (edgeR, DESeq2, and voom). Results from the simulation studies showed that BALLI correctly controlled the type-1 error rates at the various nominal significance levels, and produced better statistical power and precision estimates than those of other competing methods in various scenarios. Furthermore, BALLI was robust to variation of library size. It was also successfully applied to Holstein milk yield data, illustrating its practical value.

**Availability and Implementation:** BALLI is implemented as R package and freely available at http://healthstat.snu.ac.kr/software/balli/.

**Supplementary Information:** Supplementary data are available at *Bioinformatics* online

## INTRODUCTION

Transcriptomic profiles can improve our understanding of the phenotypic molecular basis of biological research, and many attempts have been made to identify differentially expressed genes (DEGs) by microarray analysis. However, microarray analysis often suffers from many systematic errors, such as hybridization and dye-based detection bias, hampering the detection of true DEGs (Dobbin, et al., 2005; Okoniewski and Miller, 2006). Recently, high-throughput sequencing technology has markedly improved. RNA sequencing (RNA-seq), also called whole-transcriptome shotgun sequencing, uses next-generation sequencing to quantify the abundance of transcripts with several desirable features, such as increased dynamic range and the freedom from *a priori* chosen probes (Zhao, et al., 2014). Furthermore, RNA-seq is robust against systematic errors and has therefore emerged as a successful alternative to microarray analysis (Mortazavi, et al., 2008).

RNA-seq quantifies the numbers of reads aligned to particular transcripts or genes, and various approaches have been proposed to manage the RNA-seq data (Dillies, et al., 2013). There are two different types of statistical methods: read-count-based approaches and transformation-based approaches. Read-count-based approaches assume that observed read counts follow negative binomial distribution, and generalized linear regression with a logarithm as a link function is utilized. These approaches typically assume that variances include biological and technical variances; the latter indicates variance observed among measurements of the same biological unit, and the former indicates variance between different biological units, such as different subjects or different tissues of the same subject. If technical replicates are analyzed, observed read counts from technical replicates have the same means under the same conditions. Marioni et al (2008) demonstrated that the data follow a Poisson distribution, and variances in technical replicates are expected to be the same as their means for each gene (Marioni, et al., 2008). However, if biological replicates are available, means and variances of read counts are different among different biological units. Bullard et al (2010) carefully examined such variability and concluded that the biological variances were usually larger than technical variances, supporting the presence of overdispersion (Bullard, et al., 2010). Thus, negative binomial distribution has often been utilized; edgeR (Robinson, et al., 2010) and DESeq2 (Love, et al., 2014) are such methods. Transformation-based approaches assume that the transformed read counts for each gene follow the normal distribution. For example, voom calculated proportions of read counts for each gene per subject, and the log-transformed values were then assumed to follow the normal distribution, assuming that the relative proportion of technical variances becomes smaller if the read count grows larger (Law, et al., 2014).

Negative-binomial distributions for read counts and normal distributions for log-transformation of counts per million (CPM) successfully describe distributions of RNA-seq data. However, RNA-seq is relatively expensive compared with microarray, and thus, further adjustment has been made to handle the problem of small sample size. If sample size is small, the estimated variance can have large standard errors, and thus, multiple methods that incorporate prior knowledge have been proposed. For example, variances of read counts assume to be positively related to their means, and their relationships can be estimated by comparing the means and variances of read counts for all genes. This can often be utilized to shrink variance parameters (Robinson and Smyth, 2007; Tusher, et al., 2001). edgeR and DESeq2 estimate the overall dispersion parameter for all genes and are then combined with gene-wise dispersion parameters for each gene using empirical Bayesian rules. voom assumes that the variances of log-transformed CPM (log-cpm) are functionally related to their means. Locally weighted scatterplot smoothing (LOWESS) curves between the mean and residual variances of genes are then utilized to weight variance estimates of each gene.

Existing methods shrink the variance estimate of each gene toward global variance estimates or use variance estimates based on the relationships between means and variances. Such assumptions are often very useful if the sample sizes are small. However, there are multiple factors that can distort these relationships, and if they are violated, the performance of existing approaches can be affected. For example, the quality of data is highly dependent on the preparation steps, and unexpected noise, such as noise from different storage periods or sequencing organization of samples, can occur during preparation steps. Moreover, read counts of cancer tissues are more heterogeneous than those of normal tissues, and biological variances can be affected by disease status (McCarthy, et al., 2012). Thus, their effects can distort the relationship between technical and biological variances. Multiple studies have shown that misspecified relationships can lead to substantial biases (Chavance and Escolano, 2016; Litière, et al., 2008). For example, variance estimators for random effects, which are assumed to follow a normal distribution, can be seriously biased unless they are normally distributed (Litière, et al., 2008). General approaches that are not sensitive to those problems are necessary.

In this article, we present new methods for identifying DEGs with RNA-seq data, BALLI and LLI. Statistical analyses with log-transformed read counts are often more powerful than other existing methods and are relatively insensitive to various errors (Seyednasrollah, et al., 2015; Soneson and Delorenzi, 2013). Thus, we consider the log-cpm as response variables and used linear mixed effect models to estimate technical and biological variance. Furthermore, Bartlett-adjusted likelihood ratio tests were applied to correct the small sample bias (Bartlett, 1937). By allowing model comparisons among different models, our models enable robust analyses for various scenarios. For our simulations studies, artificial RNA-seq data are generated based on real data and negative binomial distributions. Our studies showed that the proposed method performed better than existing methods. The proposed methods were applied to Holstein milk yield data at the false discovery rate (FDR)-adjusted 0.1 significance level and uniquely produced significant results. The proposed methods were implemented as an R package and are freely downloadable at http://healthstat.snu.ac.kr/software/balli/.

## MATERIAL AND METHODS

### Notations

We assumed that there were *M* different groups, and the averages of the expressed read counts for each gene were compared among these groups. For case-control studies, *M* = 2. We assumed that there were *n_m_* subjects in group *m* and denoted the total sample size by *N.* Then, we have 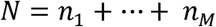. We defined dummy variables for subject *i* in group *m* by

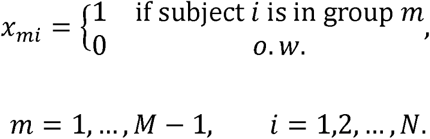

A design matrix for group variables is defined by

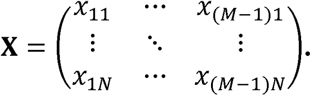

We assumed that indexes of all subjects were sorted in an ascending order of groups. Thus, the first *n*_1_ subjects were in group 1, the second *n*_2_ subjects were in group 2, and so on. We assumed that expressed read counts were observed for *G* genes and were denoted by *r_gi_* for gene g of subject *i* (*g* = 1, …, *G*). Then, the library size for subject *i, R_i_*, was equivalent to 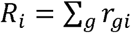. If we denoted the normalized *R_i_* with the trimmed mean of the M-value (Robinson and Oshlack, 2010) by 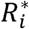, the log-transformed read counts per million of gene *g* for subject *i* were defined by:
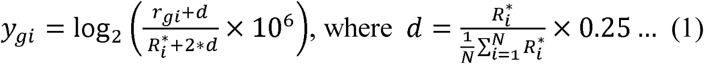

and their vector **Y*_g_*** was defined as

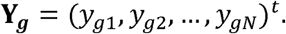

### Technical and biological variances of *y_gi_*

We assumed that *r_gi_* followed a negative binomial distribution, and its mean and variance were *μ_gi_* and 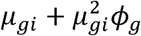, respectively. If we let the mean and variance of log_2_ *r_gi_* be *λ_gi_* and 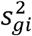, respectively, then var(*y_gi_*) can be obtained by the first order approximation as follows:
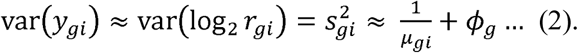

Here, 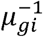 and *ϕ_g_* indicate the variances attributable to the technical and biological replicates, respectively. By the second order approximation, the technical variance, 1*/μ_gi_,* can be expressed in terms of *λ_gi_* and 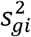 as follows:

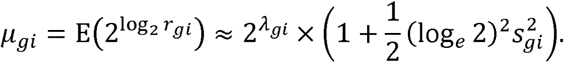

*λ_gi_* and 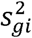 are functionally related, and both were estimated with the method used for voom-trend (Law, et al., 2014) as follows:

i. For all genes, *g* = 1,…*,G,* fit linear regressions, 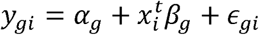, and calculate 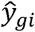. Residual variances are used as 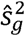. If environmental effects affect *y_gi_,* then they should be included as covariates.
ii. Calculate 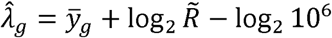 where 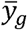 is an average of *y_gi_*, 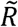 is a geometric mean of 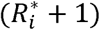 and *g* = 1, *…,G.*
iii. For 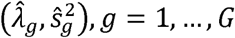, obtained from (i) and (ii), fit LOWESS curve 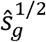 on 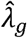.
iv. Calculate 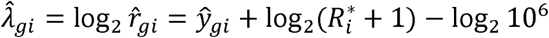 and apply LOWESS curve from (iii) to obtain 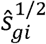.
v. Calculate 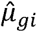 by incorporating 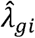 and 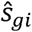 to the following equation:

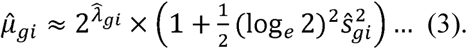

### Linear mixed effect model

We denoted a design matrix for nuisance effects including an intercept by **Z**. We let **b**_g_ and **e**_g_ be vectors for random effects and measurement errors, respectively. Denoting a *w* × *w* dimensional identity matrix by **I**_w_, we considered the following linear mixed effects model:

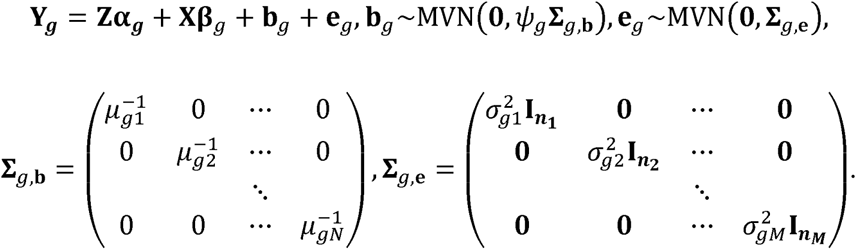

Here, *ψ_g_***Σ***_g,**b**_* and 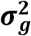 indicate technical and biological variations, respectively, according to equation (2). Notably, elements of **Σ**_*g*,b_ are obtained from equation (3), and *ψ_g_* and 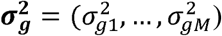 are should be estimated. Equation (2) shows that *ψ_g_* becomes 1, and we assumed that 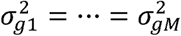. Then, our final model becomes

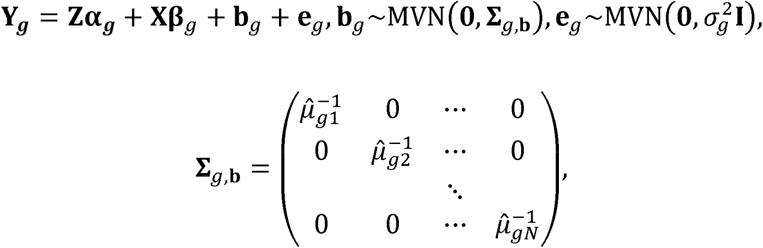

which is equivalent to

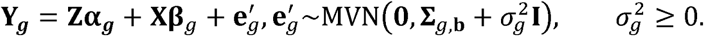

### Bartlett-adjusted profile likelihood ratio tests

Statistical analyses with RNA-seq data often use small samples, and we selected the Bartlett-adjusted likelihood ratio test for identifying DEGs. Bartlett’s adjustments make the likelihood ratio statistic close to its null distribution with reducing the order of approximation error from *O*(*N*^−1^) to *O*(*N*^−2^) and control the type-1 error rates well when the sample size is small (Bartlett, 1937). If we let **β_g_** = (β_g,i_,…, β_g,*M*−1_)*^t^*, 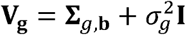 and 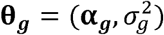, the likelihood for the proposed linear mixed model is

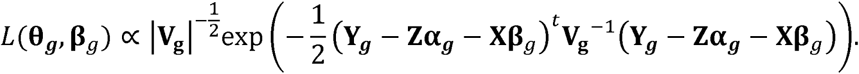

If we let 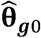 be a maximum likelihood estimate (mle) under the parameter space for the null hypothesis H_0_: **β***_g_* = 0, and 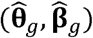 be mles of (**θ**_*g*_, **β**_*g*_) under the parameter space for null or alternative hypothesis, the likelihood ratio test for the null hypothesis H_0_: **β***_g_* = 0 can be obtained by

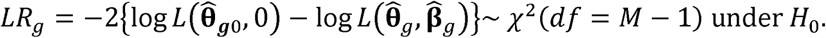

The Bartlett-adjusted likelihood ratio test 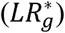 for gene *g* can be expressed by

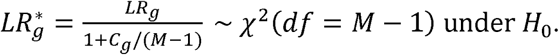

*C_g_* can be obtained based on the results of Melo et al. (2009) (Melo, et al., 2009), as follows:

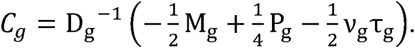

Here, D_g_, M_g_, P_g_, v_g_, and τ_g_ are scalars, and if we let **X′ = [I − Z(Z^T^V_g_^−1^Z)^−1^Z^*t*^V_g_^−1^]X** and 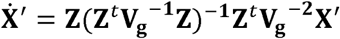, they are

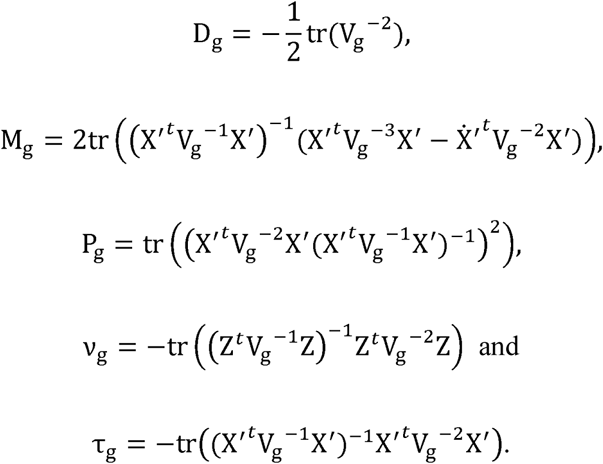

The forms of D_g_, M_g_, P_g_, v_g_, and τ_g_ depend on the structure of V_g_ and counterparts to general V_g_s are shown in the Supplementary Text A.

### Parameter estimation

The log-likelihood function for our final model is given by 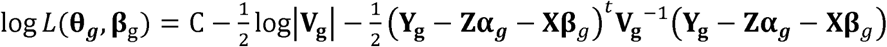, C: some constants.

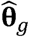 and 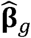 can be estimated by maximizing the log-likelihood function. Then, if we let **P = V_g_^−1^−V_g_^−1^(Z,X)((Z,X)^*t*^V_g_^−1^(Z,X))(Z,X)^*t*^V_g_^−1^**, the profile log-likelihood of 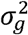 becomes

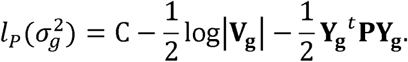

Here, **V_g_** is a function of 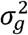, and 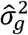 can be obtained by maximizing 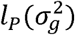 with Fisher’s Scoring method. If we let 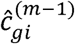 be *i*th component of 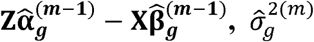 at the m step was updated by

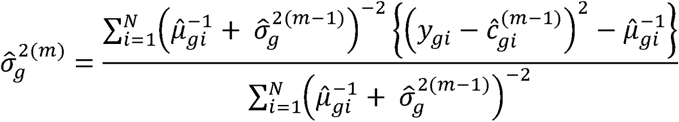

(Viechtbauer, 2007). We found that Fisher’s Scoring method was sometimes unsuccessful for estimating 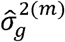, and in such cases, we used Brent’s derivative free method (Brent, 1973) with the *optimize* function in R. We assumed that 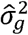 was non-negative. 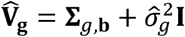, and if 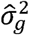 is equal to zero, 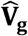 becomes **Σ***_g,**b**_*. Then, 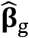 and 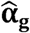 can be obtained by

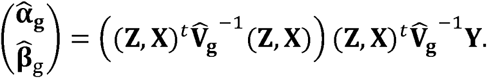

The Bartlett-adjusted likelihood ratio test requires 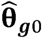, which maximizes the likelihood under the null hypothesis. Under the null hypothesis, if we let **P_0_ = V_g_^−1^** − **V_g_^−1^Z(Z^*t*^V_g_^−1^Z)Z^*t*^V_g_^−1^**, the profile log-likelihood of 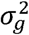 becomes

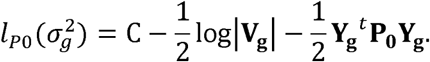

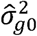 was estimated with the Fisher’s scoring method, and if we incorporated 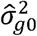 to **V_g0_** and denoted it as 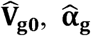 could be obtained by

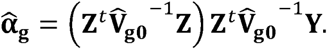

### Datasets

We considered two real datasets consisting of unrelated Nigerian and Holstein samples, respectively. Nigerian subjects were participated in the International HapMap Project and were composed of 29 males and 40 females (Pickrell, et al., 2010). The read counts were downloaded from the ReCount website (Frazee, et al., 2011). Holstein data were obtained to identify genes associated with milk yield and consisted of high and low milk yielding groups whose number of subjects is 9 and 12, respectively (Seo, et al., 2016). Furthermore, parity and lactation periods were available and were considered as covariates. Steps for transformation from the raw sequence data to read counts are shown in Supplementary Text B. Based on count data, we generated simulation data and the steps are described in the Supplementary Text C.

## RESULTS

### Simulation studies with Nigerian RNA-seq data

We applied the proposed linear mixed models to the simulated data based on Nigerian RNA-seq data and calculated empirical type-1 error rates and statistical powers with these models. The data were then compared with DESeq2 (v1.14.1), edgeR (v3.16.5) and voom (v3.30.13). Table 1, Supplementary Table 1 and Figure 1 show results from simulation studies based on Nigerian RNA-seq data. Nigerian RNA-seq data consisted of 52,580 genes, and after filtering genes whose total read counts across samples were smaller than one tenth of the sample size, each replicate had around 10,000–10,500 genes. Empirical type-1 error rates and powers were estimated with 20 replicates. Table 1 and Supplementary Table 1 assumed δ = 0, and thus, their estimates indicated the empirical type-1 error rates. For the proposed methods, we assumed that *ψ_g_* = 1 and 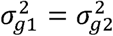, and the proposed methods with and without Bartlett’s corrections are denoted as BALLI and LLI, respectively, for the remainder of this article. According to Table 1 and Supplementary Table 1, BALLI and voom always controlled the nominal type-1 error rates correctly. LLI also successfully controlled the nominal type-1 error rates if *N* was larger than or equal to 20. However, if *N* = 12 or 16, *p* values by LLI were inflated. edgeR showed the least performance, and the estimated type-1 error rates were always inflated at 0.05, 0.01, and 0.005 nominal significance levels. Interestingly, DESeq2 tended to be conservative at 0.1 and 0.05, but liberal at 0.01 and 0.005 nominal significance levels. Thus, we could conclude that the proposed linear mixed model with Bartlett’s correction reasonably controlled the type-1 error, and Bartlett’s correction was required if the sample size was less than 20.

**Table 1.** Estimated type-1 error rates with simulation data based on Nigerian data. Estimated type-1 error rates by BALLI, DESeq2, edgeR, LLI and voom and their 95% confidence levels were estimated for *N* = 12,16,20 and 24. The type-1 error rates are marked by bold font if their 95% confidence levels include or lower than the nominal significant level *α*.

| $\alpha$ | $N = 12$ | | | | | $N = 16$ | | | | |
| --- | --- | --- | --- | --- | --- | --- | --- | --- | --- | --- |
|  | BALLI | DESeq2 | edgeR | LLI | voom | BALLI | DESeq2 | edgeR | LLI | voom |
| 0.1 | **0.10722**(<br>**0.08803**,<br>**0.12640**) | **0.09581**(<br>**0.07622**,<br>**0.11540**) | 0.12502(<br>0.10744,<br>0.14260) | 0.14288(<br>0.12020,<br>0.16556) | **0.10497**(<br>**0.08608**,<br>**0.12386**) | **0.09810**(<br>**0.07994**,<br>**0.11625**) | **0.09133**(<br>**0.07252**,<br>**0.11015**) | 0.12072(<br>0.10319,<br>0.13826) | 0.12429(<br>0.10349,<br>0.14509) | **0.10640**(<br>**0.08854**,<br>**0.12425**) |
| 0.05 | **0.05313**(<br>**0.04158**,<br>**0.06469**) | **0.05203**(<br>**0.03888**,<br>**0.06518**) | 0.06816(<br>0.05662,<br>0.07970) | 0.07703(<br>0.06158,<br>0.09247) | **0.05147**(<br>**0.03992**,<br>**0.06303**) | **0.04811**(<br>**0.03602**,<br>**0.06021**) | **0.04928**(<br>**0.03602**,<br>**0.06293**) | 0.06605(<br>0.05395,<br>0.07814) | 0.06655(<br>0.05194,<br>0.08117) | **0.05291**(<br>**0.04107**,<br>**0.06474**) |
| 0.01 | **0.01000(**<br>**0.00748,**<br>**0.01252)** | **0.01393(**<br>**0.00942,**<br>**0.01843)** | 0.01785(<br>0.01386,<br>0.02184) | 0.01917(<br>0.01441,<br>0.02392) | **0.00957(**<br>**0.00672,**<br>**0.01241)** | **0.00977(**<br>**0.00603,**<br>**0.01351)** | **0.01356(**<br>**0.00790,**<br>**0.01922)** | 0.01688(<br>0.01240,<br>0.02136) | 0.01573(<br>0.01035,<br>0.02112) | **0.01065(**<br>**0.00701,**<br>**0.01429)** |
| 0.005 | **0.00504(**<br>**0.00371,**<br>**0.00638)** | 0.00834(<br>0.00553,<br>0.01116) | 0.01063(<br>0.00811,<br>0.01314) | 0.01016(<br>0.00748,<br>0.01284) | **0.00449(**<br>**0.00303,**<br>**0.00594)** | **0.00481(**<br>**0.00273,**<br>**0.00689)** | **0.00787(**<br>**0.00410,**<br>**0.01164)** | 0.00974(<br>0.00689,<br>0.01260) | 0.00862(<br>0.00527,<br>0.01198) | **0.00531(**<br>**0.00323,**<br>**0.00740)** |
| $\alpha$ | $N = 20$ | | | | | $N = 24$ | | | | |
|  | BALLI | DESeq2 | edgeR | LLI | voom | BALLI | DESeq2 | edgeR | LLI | voom |
| 0.1 | **0.09292(**<br>**0.07721,**<br>**0.10863)** | **0.09016(**<br>**0.07371,**<br>**0.10662)** | 0.11795(<br>0.10291,<br>0.13299) | **0.11330(**<br>**0.09560,**<br>**0.13100)** | **0.10581(**<br>**0.08793,**<br>**0.12369)** | **0.09300(**<br>**0.07477,**<br>**0.11123)** | **0.08930(**<br>**0.06998,**<br>**0.10862)** | **0.11535(**<br>**0.09727,**<br>**0.13344)** | **0.10886(**<br>**0.08888,**<br>**0.12883)** | **0.09738(**<br>**0.07824,**<br>**0.11651)** |
| 0.05 | **0.04441(**<br>**0.03436,**<br>**0.05446)** | **0.04736(**<br>**0.03597,**<br>**0.05875)** | 0.06353(<br>0.05333,<br>0.07372) | **0.05834(**<br>**0.04603,**<br>**0.07065)** | **0.05343(**<br>**0.04226,**<br>**0.06459)** | **0.04528(**<br>**0.03330,**<br>**0.05726)** | **0.04822(**<br>**0.03470,**<br>**0.06174)** | 0.06513(<br>0.05302,<br>0.07725) | **0.05621(**<br>**0.04247,**<br>**0.06995)** | **0.04771(**<br>**0.03541,**<br>**0.06002)** |
| 0.01 | **0.00797(**<br>**0.00537,**<br>**0.01058)** | **0.01156(**<br>**0.00750,**<br>**0.01563)** | 0.01541(<br>0.01212,<br>0.01870) | **0.01221(**<br>**0.00858,**<br>**0.01583)** | **0.00949(**<br>**0.00699,**<br>**0.01200)** | **0.00846(**<br>**0.00512,**<br>**0.01181)** | **0.01269(**<br>**0.00739,**<br>**0.01800)** | 0.01681(<br>0.01244,<br>0.02117) | **0.01224(**<br>**0.00762,**<br>**0.01685)** | **0.00921(**<br>**0.00570,**<br>**0.01272)** |
| 0.005 | **0.00367(**<br>**0.00231,**<br>**0.00504)** | **0.00632(**<br>**0.00378,**<br>**0.00885)** | 0.00873(<br>0.00677,<br>0.01068) | **0.00626(**<br>**0.00414,**<br>**0.00838)** | **0.00443(**<br>**0.00311,**<br>**0.00575)** | **0.00419(**<br>**0.00236,**<br>**0.00602)** | **0.00717(**<br>**0.00385,**<br>**0.01050)** | 0.00962(<br>0.00690,<br>0.01235) | **0.00625(**<br>**0.00368,**<br>**0.00883)** | **0.00432(**<br>**0.00248,**<br>**0.00617)** |

**Figure 1.**
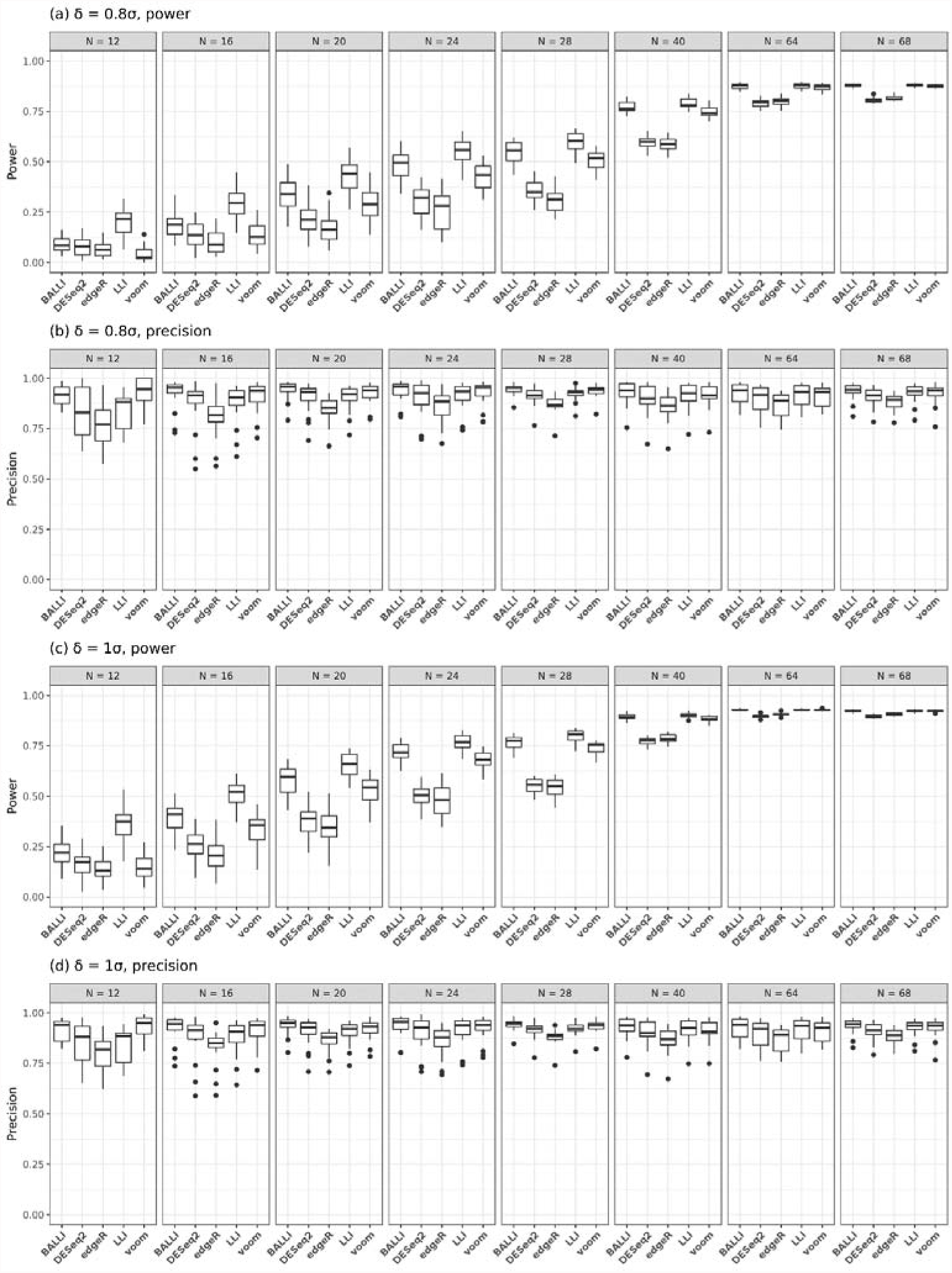
Estimated Powers and Precisions with simulation data based on Nigerian RNA-seq data. Statistical powers of BALLI, DESeq2, edgeR, LLI and voom were estimated at FDR-adjusted 0.1 significance level when δ = 0.8σ, 1σ and

Figure 1 shows estimated powers and precisions at the FDR-adjusted 0.1 significance level when δ = 0.8σ or 1σ and N = 12, 16, 20, 24, 28, 40, 64, or 68. Figure 1a and 1c show the statistical power estimates, and Figure 1b and 1d show the precision. Precision indicates the proportions of DEGs among genes for which FDR-adjusted *p* values are less than 0.1. According to Figure 1a and 1c, LLI outperformed other methods in terms of power (Figure 1a and 1c). However, it should be noted that this method showed worse precision than BALLI and voom if *N* was less than 20 (Figure 1b and 1d), suggesting that LLI had larger falsepositive rates than BALLI and voom when *N* was less than 20. The precision of LLI was increased if *N* was sufficiently large. In terms of both power and precision, the best performance was always obtained by BALLI. For example, when *N* = 20 and δ = σ, the estimated power of BALLI was 0.577, followed by voom (0.526) and DESeq2 (0.376). The estimated precision of BALLI was 0.936, and those of voom and DESeq2 were 0.919 and 0.906, respectively. If *N* = 16 and δ = 0.8σ, the estimated power and precision of BALLI were 0.188 and 0.926, which were higher than those of DESeq2 (0.137, 0.870) and voom (0.138, 0.910).

### Simulation studies with randomly generated RNA-seq data

RNA-seq data are generally known to follow the negative binomial distribution, and we conducted simulation studies with RNA-seq data generated from negative binomial distributions. First, we assumed that library sizes were the same among subjects. The overall trend of the estimated type-1 error rate was similar to that of simulation studies based on Nigerian RNA-seq data. Estimated type-1 error rates by voom and BALLI usually maintained the nominal significance levels (Table 2 and Supplementary Table 2). *P* values obtained from LLI and edgeR tended to be inflated, but the amount of inflation by LLI was small compared with that of edgeR. DESeq2 generally showed deflation of type-1 error rates at 0.1 and 0.05 nominal significance levels. Second, we considered the effects of library size variance on statistical analyses. Data with unequal library sizes among subjects were generated by negative binomial distribution whose mean parameters (*a_gi_*) were the product of mean estimates, under the equal library size assumption, and random numbers from *U*(*u,* 2 − *u*), where *u* = 0.2,0.4,0.6,0.8 or 1, and dispersion parameters were estimated from 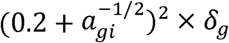, where 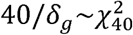. If *u* became smaller, the library size had larger variances. Figure 2 shows the estimated type-1 error rates at the 0.05 significance level according to different choices of u. Figure 2 shows that voom was sensitive to the amount of library size variation and became conservative in the context of large library size variation. Compared with voom, BALLI and LLI were robust with regard to u. The estimated type-1 error rates of LLI were affected by sample size. If *N* was larger than or equal to 40, LLI controlled the type-1 error rates the most correctly and was not affected by the library size variation. BALLI was slightly conservative, but the amount remained constant. Results at the 0.005 significance level are provided in Supplementary Figure 1, and the general pattern was similar to that in Figure 2, except that DESeq2 was relatively closer to the base line (Supplementary Figure 1). Therefore, we could conclude that the performances of BALLI and LLI were robust, and we recommend using BALLI if 10 ≤ N ≤ 40 and LLI if N > 40.

**Table 2.** Estimated type-1 error rates with simulation data based on simulated RNA-seq data from negative binomial distribution. Estimated type-1 error rates by BALLI, DESeq2, edgeR, LLI and voom and their 95% confidence levels were estimated for *N* = 12,16,20 and 24. The type-1 error rates are marked by bold font if their 95% confidence levels include or lower than the nominal significant level *α*.

| $\alpha$ | $N = 12$ | | | | | $N = 16$ | | | | |
| --- | --- | --- | --- | --- | --- | --- | --- | --- | --- | --- |
|  | BALLI | DESeq2 | edgeR | LLI | voom | BALLI | DESeq2 | edgeR | LLI | Voom |
| 0.1 | **0.09728**<br>(0.09619, 0.09838) | **0.08127**<br>(0.08013, 0.08241) | 0.12351<br>(0.12234, 0.12468) | 0.13228<br>(0.13118, 0.13339) | **0.09869**<br>(0.09750, 0.09989) | **0.09214**<br>(0.09054, 0.09374) | **0.08275**<br>(0.08145, 0.08405) | 0.12219<br>(0.12094, 0.12345) | 0.11895<br>(0.11711, 0.12079) | **0.10026**<br>(0.09872, 0.10180) |
| 0.05 | **0.04792**<br>(0.04708, 0.04876) | **0.04082**<br>(0.03997, 0.04168) | 0.05743<br>(0.05644, 0.05842) | 0.06983<br>(0.06877, 0.07089) | **0.04915**<br>(0.04806, 0.05023) | **0.04607**<br>(0.04536, 0.04677) | **0.04261**<br>(0.04163, 0.04359) | 0.05859<br>(0.05744, 0.05974) | 0.06435<br>(0.06333, 0.06536) | **0.05046**<br>(0.04950, 0.05142) |
| 0.01 | **0.00990**<br>(0.00943, 0.01037) | **0.00877**<br>(0.00839, 0.00915) | 0.01272<br>(0.01227, 0.01317) | 0.01798<br>(0.01743, 0.01853) | **0.00997**<br>(0.00948, 0.01046) | **0.00955**<br>(0.00911, 0.00999) | **0.00980**<br>(0.00934, 0.01027) | 0.01379<br>(0.01332, 0.01427) | 0.01575<br>(0.01525, 0.01625) | **0.01032**<br>(0.00992, 0.01073) |
| 0.005 | **0.00524**<br>(0.00488, 0.00559) | **0.00497**<br>(0.00463, 0.00530) | 0.00703<br>(0.00668, 0.00737) | 0.01034<br>(0.00986, 0.01082) | **0.00522**<br>(0.00489, 0.00555) | **0.00480**<br>(0.00453, 0.00506) | 0.00550<br>(0.00523, 0.00576) | 0.00757<br>(0.00726, 0.00788) | 0.00857<br>(0.00816, 0.00899) | **0.00517**<br>(0.00490, 0.00544) |
| $\alpha$ | $N = 20$ | | | | | $N = 24$ | | | | |
|  | BALLI | DESeq2 | edgeR | LLI | voom | BALLI | DESeq2 | edgeR | LLI | voom |
| 0.1 | **0.09188** | **0.08277** | 0.11842 | 0.11062 | **0.09941** | **0.09491** | **0.08516** | 0.11899 | 0.11056 | **0.10126** |
|  | (0.09069,<br>0.09307) | (0.08179,<br>0.08375) | (0.11718,<br>0.11967) | (0.10960,<br>0.11164) | (0.09834,<br>0.10048) | (0.09349,<br>0.09634) | (0.08383,<br>0.08650) | (0.11755,<br>0.12043) | (0.10909,<br>0.11203) | (0.09972,<br>0.10280) |
| 0.05 | 0.04421<br>(0.04335,<br>0.04506) | 0.04184<br>(0.04088,<br>0.04280) | 0.05927<br>(0.05780,<br>0.06075) | 0.05732<br>(0.05625,<br>0.05840) | 0.04942<br>(0.04853,<br>0.05031) | 0.04585<br>(0.04474,<br>0.04696) | 0.04335<br>(0.04233,<br>0.04437) | 0.06308<br>(0.06207,<br>0.06409) | 0.05684<br>(0.05568,<br>0.05799) | 0.05021<br>(0.04936,<br>0.05105) |
| 0.01 | 0.00887<br>(0.00852,<br>0.00922) | 0.00967<br>(0.00913,<br>0.01021) | 0.01344<br>(0.01299,<br>0.01389) | 0.01326<br>(0.01295,<br>0.01356) | 0.01005<br>(0.00970,<br>0.01040) | 0.00897<br>(0.00849,<br>0.00944) | 0.00993<br>(0.00955,<br>0.01032) | 0.01377<br>(0.01328,<br>0.01426) | 0.01265<br>(0.01217,<br>0.01313) | 0.01007<br>(0.00963,<br>0.01051) |
| 0.005 | 0.00455<br>(0.00432,<br>0.00479) | 0.00536<br>(0.00504,<br>0.00568) | 0.00738<br>(0.00698,<br>0.00778) | 0.00718<br>(0.00687,<br>0.00748) | 0.00503<br>(0.00477,<br>0.00529) | 0.00440<br>(0.00419,<br>0.00462) | 0.00515<br>(0.00484,<br>0.00547) | 0.00733<br>(0.00696,<br>0.00770) | 0.00678<br>(0.00640,<br>0.00716) | 0.00498<br>(0.00466,<br>0.00530) |

**Figure 2.**
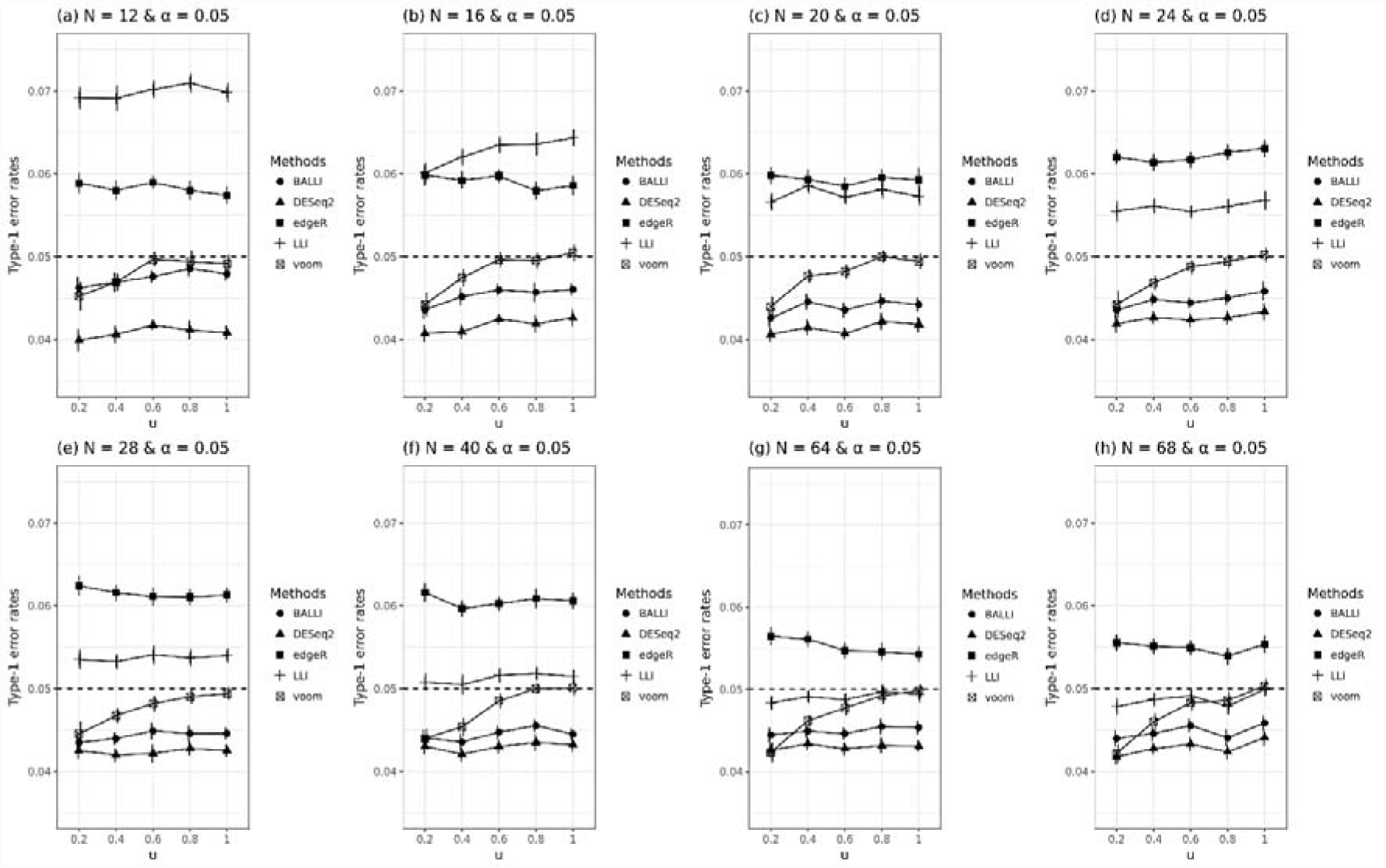
Effect of varying library sizes on the type-1 error rates. Type-1 error rates were estimated by BALLI, DESeq2, edgeR, LLI and voom when u = 0.2, 0.4, 0.6, 0.8 and 1 and sample size (N) is 12, 16, 20, 24, 28, 40, 64 or 68 at the 0.05 nominal significance level.

Figure 3 and Supplementary Figure 2 show the estimated statistical powers and precision according to different choices of u. BALLI usually had the best estimated power and precision, as was observed in simulation studies based on Nigerian RNA-seq data. For example, when *u* = 0.2, *N* = 20, and δ = 0.8σ, the estimated power by BALLI was 0.641, whereas those for DESeq2 and voom were 0.480 and 0.571, respectively (Figure 3a). Results when *u* = 0.2 and δ = 1σ in Figure 3c are very similar as those for Figure 3a. Figure 3b and Figure 3d also shows that BALLI achieve the best estimated precisions. Similar patterns were observed when *u* = 0.4, 0.6, 0.8, and 1 (Supplementary Figure 2). In summary, we can conclude that BALLI shows better performance than other methods.

**Figure 3.**
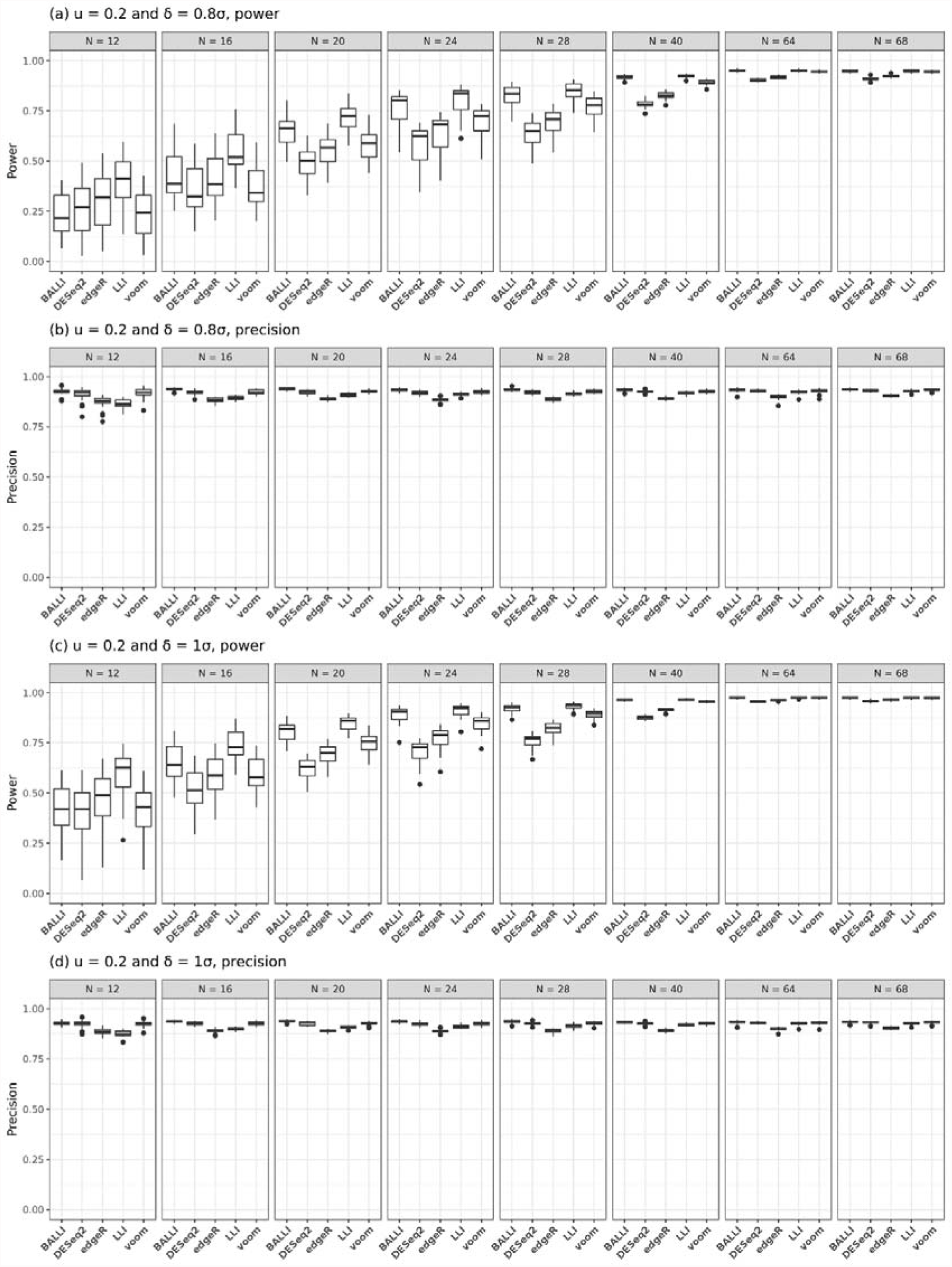
Effect of varying library sizes on the statistical power and precision. Statistical powers and precision for BALLI, DESeq2, edgeR, LLI and voom were empirically estimated at FDR-adjusted 0.1 significance level when u = 0.2, δ = 0.8σ or 1σ and sample size (N) is 12, 16, 20, 24, 28, 40, 64 or 68.

### DEGs of Holstein milk data

Holstein milk data, consisting of 21 Holstein cows, were generated to detect genes related to the productivity of daily milk. High and low milk yields were considered the primary exposure variables, and parity and lactation period were included as covariates (Seo, et al., 2016). In this study, twelve tentative DEGs were chosen, and technically validated using quantitative real time polymerase chain reaction (qRT-PCR). qRT-PCR was conducted with QuantiTect SYBR Green RT-PCR Master Mix (Qiagen, Valencia, CA, USA), and a 7500 Fast Sequence Detection System (Applied Biosystems, Foster City, CA, USA) was used to confirm whether the twelve tentative genes were true DEGs. Among the twelve genes, nine (*TOX4, HNRNPL, SPTSSB, NOS3, C25H16orf88, KALRN, SLC4A1, NLN,* and *PMCH*) were significantly validated. According to Seo et al (2016), however, no DEGs including the nine genes were found at FDR 0.1 significance level by DESeq2 and voom as well as their methods due to the lack of statistical power (Seo, et al., 2016). Our proposed methods and existing methods (DESeq2, edgeR, and voom) were applied to the data analysis. LLI only detected significant differences for the *TOX4* gene between the high and low milk yield groups at the FDR-adjusted 0.1 significance level, but other methods did not detect any significant genes. The FDR-adjusted *p* value of *TOX4* by BALLI was 0.1272, which was much smaller than those of DESeq2, edgeR, and voom. Table 3 shows *p* values for the nine genes, including *TOX4. P* values for the nine genes obtained by LLI and BALLI were small compared with those obtained from other methods. We also analyzed all genes with proposed methods; Figure 4 shows the number of genes that were significant at the 0.001 nominal significance level. There were no DEGs that were commonly significant only for all existing methods (DESeq2, edgeR and voom). Four genes, including *HNRNPL,* were detected as DEGs by only BALLI and LLI (Figure 4). Table 4 shows eight genes that were commonly significant by BALLI, DESeq2, edgeR, LLI, and voom at the 0.005 nominal significance level. Of the eight genes, all genes had the lowest *p* values in LLI, and three genes had lower *p* values in BALLI than in DESeq2, edgeR, and voom. Simulation studies revealed that LLI tended to be liberal, and the results may be inflated. However, BALLI controlled the nominal significance level, and *p* values by BALLI were expected to be statistically valid. Therefore, we concluded that the proposed method, BALLI, worked well for real data analysis.

**Table 3.**
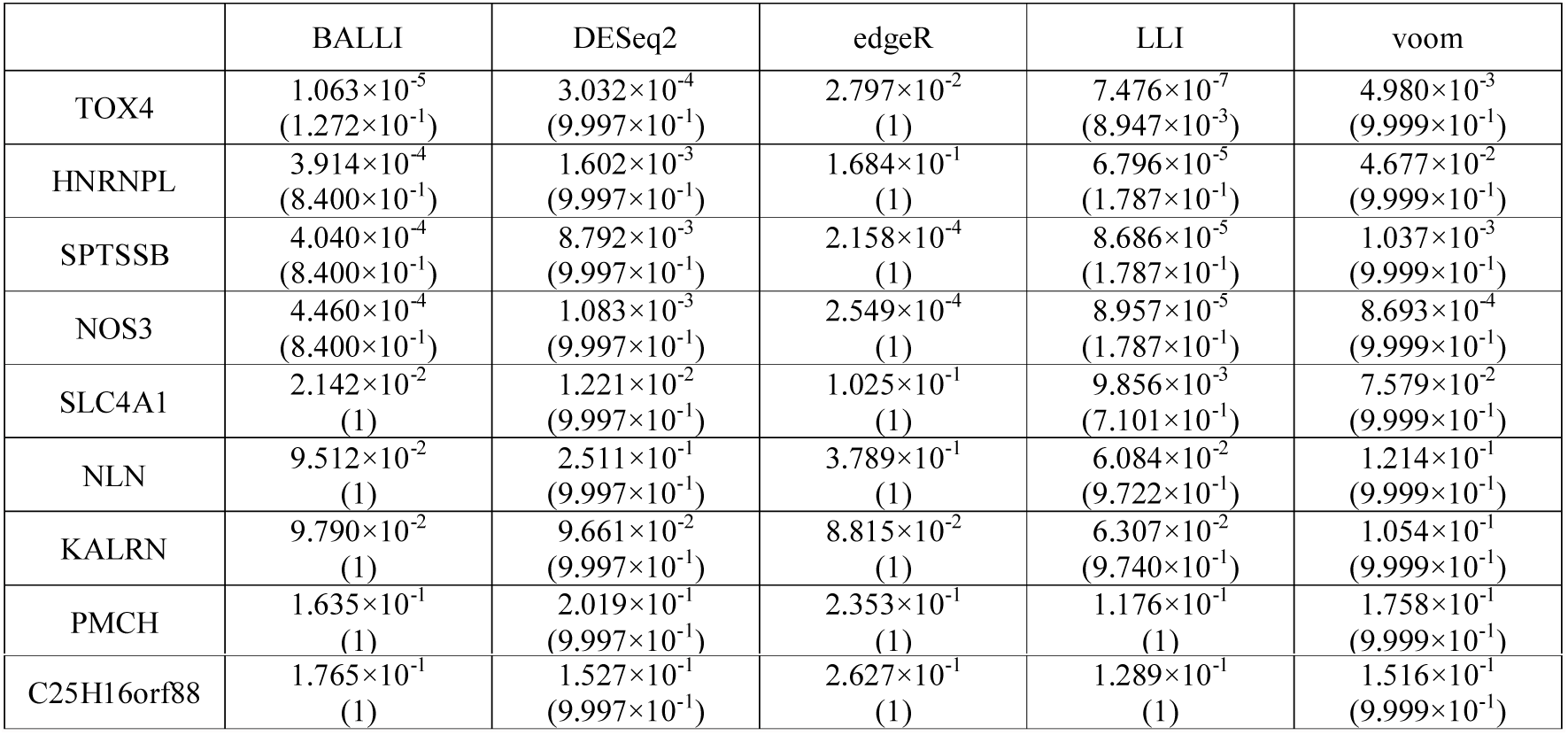
True DEG analysis results of Holstein milk data. Holstein milk data was analyzed by BALLI, DESeq2, edgeR, LLI and voom and their p values (FDRs) are provided

**Figure 4.**
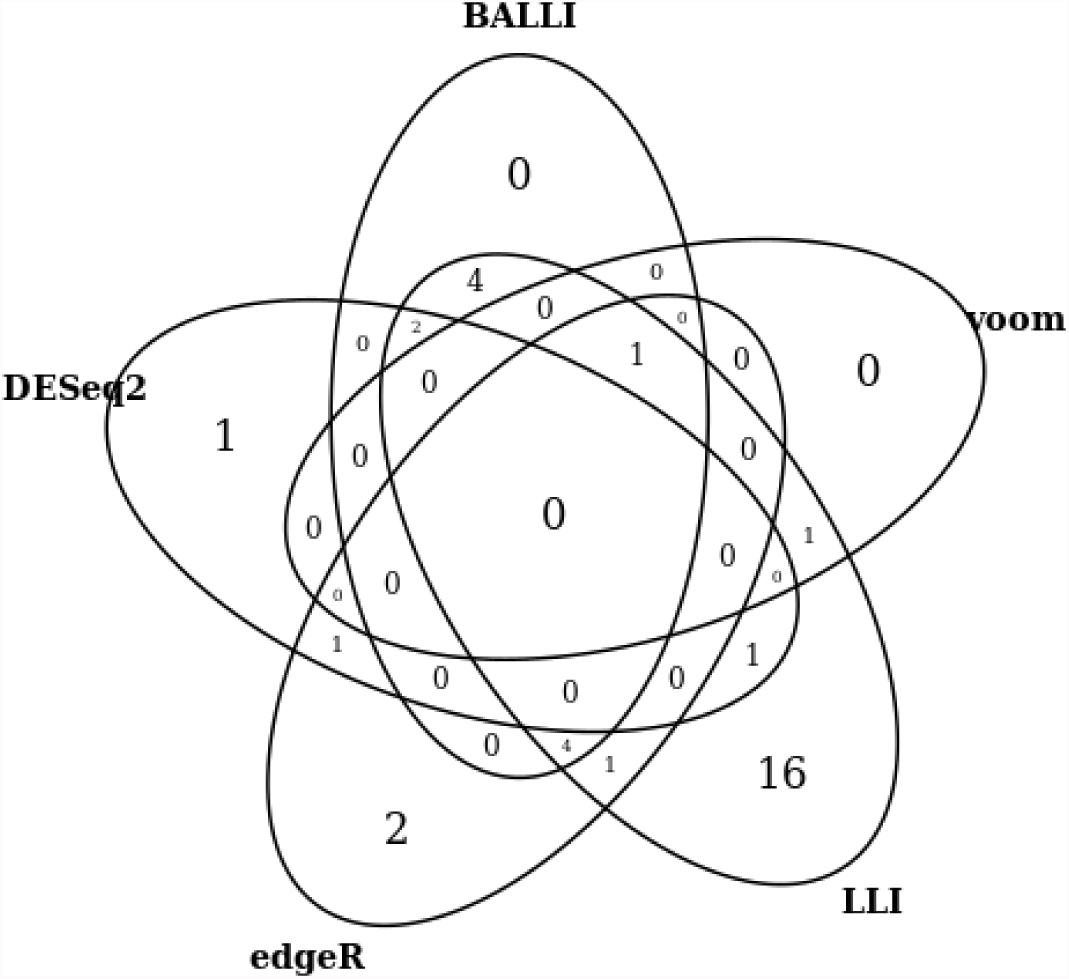
Significant genes of Holstein milk data. Venn diagram was provided with significant genes at the 0.001 nominal significance level by BALLI, DESeq2, edgeR, LLI and voom

**Table 4.** Significant genes in all methods of Holstein milk data. Gene lists of Holstein milk data siginificant in nominal 0.005 significant level for all methods (BALLI, DESeq2, edgeR, LLI and voom) and their p values (FDRs) are provided.

|  | BALLI | DESeq2 | edgeR | LLI | voom |
| --- | --- | --- | --- | --- | --- |
| NOS3 | $4.460 \times 10^{-4}$<br>(8.400 $\times 10^{-1}$) | $1.083 \times 10^{-3}$<br>(9.997 $\times 10^{-1}$) | $2.549 \times 10^{-4}$<br>(1) | $8.957 \times 10^{-5}$<br>(1.787 $\times 10^{-1}$) | $8.693 \times 10^{-4}$<br>(9.999 $\times 10^{-1}$) |
| SPESP1 | $1.065 \times 10^{-3}$<br>(8.570 $\times 10^{-1}$) | $4.901 \times 10^{-3}$<br>(9.997 $\times 10^{-1}$) | $1.669 \times 10^{-3}$<br>(1) | $2.579 \times 10^{-4}$<br>(2.385 $\times 10^{-1}$) | $4.001 \times 10^{-3}$<br>(9.999 $\times 10^{-1}$) |
| CHST1 | $1.330 \times 10^{-3}$<br>(8.570 $\times 10^{-1}$) | $3.074 \times 10^{-3}$<br>(9.997 $\times 10^{-1}$) | $1.872 \times 10^{-3}$<br>(1) | $3.166 \times 10^{-4}$<br>(2.385 $\times 10^{-1}$) | $6.108 \times 10^{-4}$<br>(9.999 $\times 10^{-1}$) |
| LEPREL1 | $1.369 \times 10^{-3}$<br>(8.570 $\times 10^{-1}$) | $3.775 \times 10^{-3}$<br>(9.997 $\times 10^{-1}$) | $3.297 \times 10^{-3}$<br>(1) | $3.334 \times 10^{-4}$<br>(2.385 $\times 10^{-1}$) | $1.487 \times 10^{-3}$<br>(9.999 $\times 10^{-1}$) |
| JUB | $1.447 \times 10^{-3}$<br>(8.570 $\times 10^{-1}$) | $2.559 \times 10^{-3}$<br>(9.997 $\times 10^{-1}$) | $2.445 \times 10^{-3}$<br>(1) | $3.674 \times 10^{-4}$<br>(2.385 $\times 10^{-1}$) | $2.804 \times 10^{-3}$<br>(9.999 $\times 10^{-1}$) |
| MIA | $1.504 \times 10^{-3}$<br>(8.570 $\times 10^{-1}$) | $3.792 \times 10^{-4}$<br>(9.997 $\times 10^{-1}$) | $2.247 \times 10^{-3}$<br>(1) | $3.666 \times 10^{-4}$<br>(2.385 $\times 10^{-1}$) | $2.432 \times 10^{-3}$<br>(9.999 $\times 10^{-1}$) |
| CLDN6 | $2.646 \times 10^{-3}$<br>(1) | $2.096 \times 10^{-3}$<br>(9.997 $\times 10^{-1}$) | $2.414 \times 10^{-3}$<br>(1) | $7.377 \times 10^{-4}$<br>(3.270 $\times 10^{-1}$) | $1.542 \times 10^{-3}$<br>(9.999 $\times 10^{-1}$) |
| PALMD | $3.733 \times 10^{-3}$<br>(1) | $1.676 \times 10^{-3}$<br>($9.997 \times 10^{-1}$) | $2.648 \times 10^{-3}$<br>(1) | $1.127 \times 10^{-3}$<br>($3.700 \times 10^{-1}$) | $3.032 \times 10^{-3}$<br>($9.999 \times 10^{-1}$) |

## DISCUSSION

In this article, we suggested new methods, designated BALLI and LLI, for identifying DEGs with RNA-seq data. We assumed that log-cpm values of read counts asymptotically followed normal distributions, and the linear mixed effects model with Bartlett’s correction was proposed. The proposed methods were compared with existing methods, such as DESeq2, edgeR, and voom, with extensive simulation studies. According to our results, negative-binomial-based approaches often failed to preserve the nominal type-1 error rates. For example, *p* values from edgeR were inflated. DESeq2 tended to be conservative and suffered from large false-negative rates. However, the proposed method with Bartlett’s correction, BALLI, preserved the nominal type-1 error rates and was the most powerful method other than LLI. Unless sample sizes were small, LLI controlled the type-1 error rates as well and was the most powerful method. Therefore, we recommend using LLI if the sample size is sufficiently large (e.g., larger than 40); otherwise, it is better to use BALLI.

Furthermore, we evaluated the effects of library size variations on statistical analyses. We found that library size variance could affect the estimated type-1 error rates, and the effect was the largest for voom. Library sizes are affected by multiple factors, such as the amount of mRNA and the sequencing instrument, which can generate substantial variation among library sizes for subjects. Our simulation studies showed that BALLI was robust with regard to library size variation in samples of various sizes and was a reasonable choice if large library size variance was observed.

The proposed methods assumed that log-cpm values of read counts asymptotically followed a normal distribution and that their variances were approximately equal to 1/*μ* + ϕ with the first order approximation. In addition, voom considered log-cpm value as a response and assumed that they were normally distributed. However, our simulation studies revealed the superiority of the proposed methods compared with voom, which was found to be attributable to their different variance structures. For the proposed methods, 1/*μ* + ϕ was derived from the first-order approximation of the negative binomial distribution and thus may be a natural assumption for RNA-seq data. Furthermore, for 1/*μ* + ϕ, ϕ obviously indicates the overdispersion parameter, and biological and technical variances can be estimated with BALLI. However, voom assumes ϕ /*μ*, and the amount attributable to biological or technical variances cannot be clearly defined.

We also suggested the most flexible and general linear mixed model for log-cpm. The proposed model assumed that the variance of log-cpm was ϕ/*μ* + ϕ and had the most generalized variance parameter space. Incorporation of ϕ = 1 yielded BALLI and LLI, and ϕ = 0 yielded voom. We found that BALLI was the most efficient in the considered scenarios; however, in real data analyses, various factors affected variance structure. For example, subjects with different ethnicities can cause ϕ to be larger than 1, and thus, a better model may differ according to RNA-seq data. ϕ and ϕ can be estimated with the proposed linear mixed model by implementing only a simple modification, and thus, we can choose the best model using AIC or likelihood ratio tests. The selected models can then be utilized to identify DEGs. This model was implemented as an R package and can be downloaded from http://healthstat.snu.ac.kr/software/balli/. Furthermore, the proposed methods can be easily extended to various scenarios via a simple modification. For example, repeatedly observed data or multivariate phenotypes can be analyzed by adding some random effects. Maximizing the likelihood for negative binomial distributions with random effects is computationally intensive, but the proposed methods can easily obtain variance parameter estimates using existing R packages, such as lme4 and nlme.

With simulation studies for various scenarios, we showed that the proposed methods were usually the most efficient. However, results from simulation studies obviously depended on various factors. Our results were obtained from simulation data based on Nigerian RNA-seq data and random samples from negative binomial distributions, but any systematic differences in RNA-seq data could generate different results, depending on sequencing errors or differences in preparation steps. Multiple studies have revealed some possible differences in these relationships, and our conclusions based on simulation studies could be limited to the considered scenarios. However, despite such limitations, we believe that our results illustrate the practical value of the proposed methods. Further studies are needed to confirm our findings and expand on the work presented herein.

## FUNDING

This research was supported by a grant of the Korea Health Technology R&D Project through the Korea Health Industry Development Institute (KHIDI), funded by the Ministry of Health & Welfare, Republic of Korea (H I15C2165).

